# GlycoMeSH: linking glycan structures to biomedical context for systematic enrichment analysis

**DOI:** 10.64898/2026.08.18.745318

**Authors:** Akihiro Kitani, Bingyuan Zhang, Koichi Himori, Yusuke Matsui

## Abstract

Glycan identification has advanced, but glycan structures remain difficult to translate into reproducible biomedical context because reusable glycan-level annotations are sparse. We present GlycoMeSH, a resource that links glycans to Medical Subject Headings (MeSH) through an inference model, a traceable association database and a glycan-set enrichment workflow. GlycoMeSH-BERT recovered ∼60% of literature-derived associations at recall@30 and expanded open-vocabulary MeSH coverage beyond closed-label baselines, without higher per-prediction accuracy. At matched candidate counts, its predictions showed motif-level semantic agreement comparable to those baselines, independently of the training labels. GlycoMeSH-DB contains 789,627 associations between 26,954 glycans and 20,302 MeSH terms. GlycoMeSH-EA returned enriched MeSH terms for glycan sets from glycomics and glycoproteomics datasets. Each association represents a biomedical context rather than a validated mechanism, and retains its source PMID or prediction score for audit. GlycoMeSH supplies the missing, evidence-traceable annotation layer that makes glycan sets directly analyzable by enrichment across glycoscience datasets.

## Introduction

Protein glycosylation regulates cell–cell interactions, immune recognition and host– pathogen interactions^1,2^. Different glycans on the same protein can alter receptor binding, stability, circulatory half-life, immunogenicity and signaling^2–5^. Their interpretation is complicated by branching, linkage and site-specific heterogeneity, through which closely related structures can mark different biological settings. Yet glycomics and glycoproteomics often end with structural descriptions because individual glycan structures are not systematically connected to a controlled, reusable vocabulary of biomedical context. This leaves a growing catalogue of identified structures without a corresponding layer for systematic interpretation.

Transcriptomic and proteomic datasets can be interpreted reproducibly with over-representation analysis (ORA), gene set enrichment analysis and curated knowledge bases^6^. Mass spectrometry and computational methods have markedly improved glycosite and glycan identification^7–10^, but identification alone does not provide the annotation layer required for analogous glycan-set analysis. In other omics fields, reusable identifiers and controlled annotations make the mapping from a measured feature to an enrichment term explicit and auditable. No equivalently broad mapping has been available for individual glycans. Consequently, detected glycans are usually summarized by composition, class or motif and interpreted through expert knowledge rather than a queryable system. Such interpretation is valuable, but it is difficult to reproduce consistently across datasets.

The bottleneck is therefore not annotation sparsity alone, but the absence of a standard mapping from glycan structure to biomedical meaning. Gene- and protein-based proxies cannot fully resolve this problem because glycan diversity is not directly encoded by genes and site-specific glycoforms can differ on the same protein. Collapsing glycoforms to their carrier proteins can therefore erase the molecular unit that differs between conditions. Without a glycan-level annotation layer, structure-to-context mappings remain implicit, difficult to reuse across studies and biased toward familiar motifs. The field consequently lacks both a shared unit of interpretation and a statistical route for asking which biomedical contexts are over-represented in a measured glycan set.

Medical Subject Headings (MeSH) offer a controlled vocabulary spanning diseases, phenotypes, molecular entities and biological processes. Because MeSH terms annotate publications rather than a particular molecular class, they provide a practical route from glycan-associated literature to reusable biomedical-context annotations, although the resulting links are document-level contexts rather than mechanistic glycan functions. Direct mapping was nevertheless limited to 27,563 associations between 3,061 glycans and 2,174 MeSH terms after filtering. These annotations covered only 4.4% of the 69,554 glycans with available IUPAC-condensed structures, leaving most structurally defined glycans outside a usable annotation space. We developed GlycoMeSH to expand these sparse links with GlycoMeSH-BERT, combine literature and predicted associations in GlycoMeSH-DB, and analyze glycan sets with GlycoMeSH-EA. The resource retains the provenance or prediction score of each association so that inference does not obscure evidence source. We evaluated held-out label recovery, open-vocabulary coverage and motif-level semantic agreement, then illustrated the resource in glycomics and glycoproteomics datasets. Our aim was to supply this missing annotation layer as a resource broad and traceable enough for glycan-set interpretation, rather than to validate individual inferred mechanisms.

## Results

A glycan-centric biomedical-context annotation and enrichment framework GlycoMeSH addresses the absence of a reusable structure-to-context annotation layer for glycan-set analysis (Fig. 1). Gene-centered resources such as Gene Ontology, KEGG and Reactome support enrichment analysis^18–20^, whereas existing glycoinformatics resources primarily provide structural, ontological, enzymatic, interaction or lectin-binding information (Table 1). GlycoMeSH complements these resources by linking glycan structures to MeSH-based biomedical contexts and making the resulting annotations directly queryable by enrichment analysis (Fig. 1a).

**Fig. 1:**
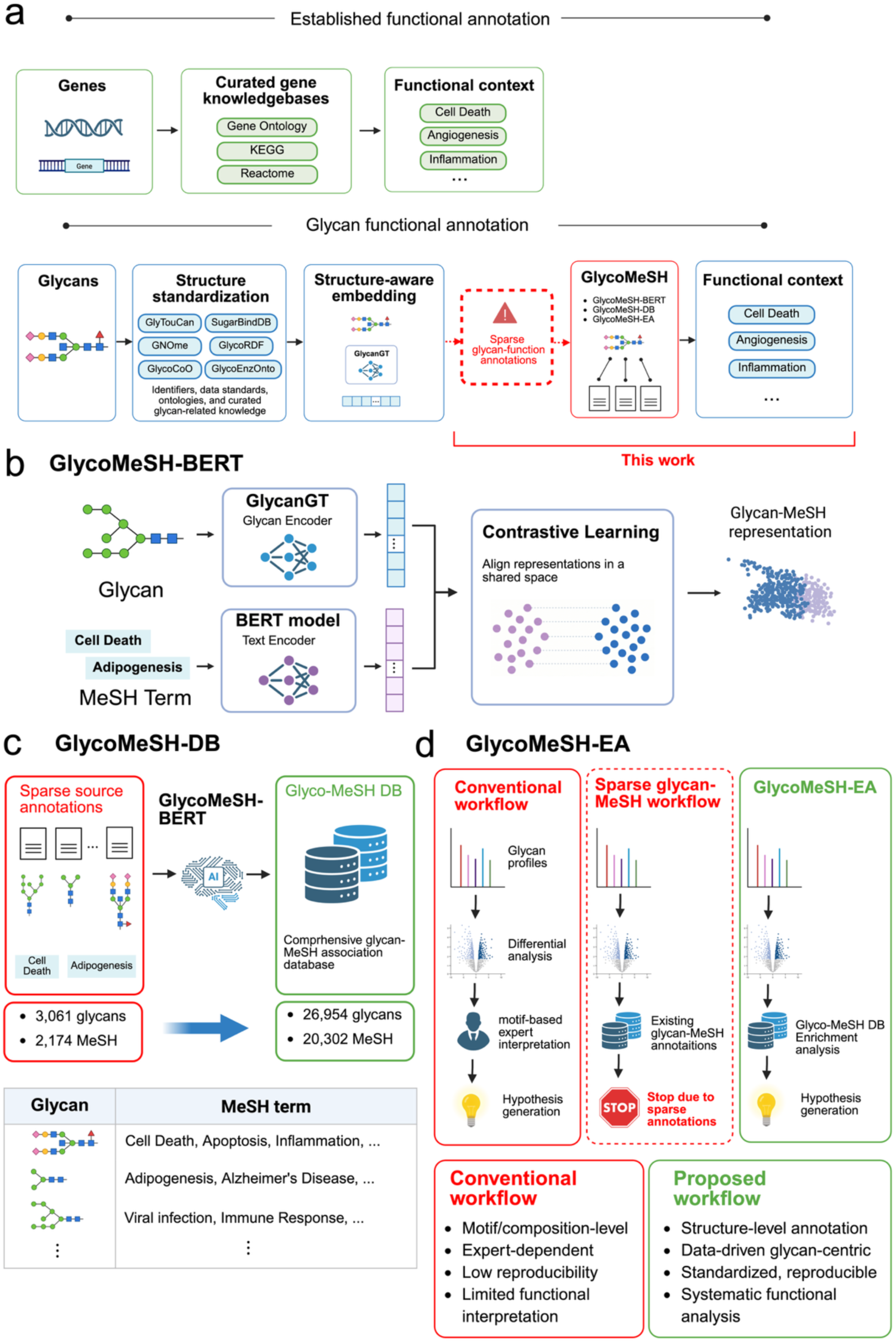
Overview of the GlycoMeSH structure-to-biomedical-context annotation and enrichment framework. a, Comparison of established gene-centered annotation resources with the remaining gap in glycan-level biomedical-context annotation. b, Construction of GlycoMeSH-BERT. Glycans and MeSH terms are embedded using pretrained models and aligned in a shared representation space by contrastive learning. c, Construction of GlycoMeSH-DB. Model-predicted associations are integrated with literature-derived annotations to expand glycan– MeSH coverage; the predicted entries are score-thresholded contexts, not individually validated knowledge. d, GlycoMeSH-EA queries GlycoMeSH-DB for glycan-set enrichment while retaining source-PMID or prediction-score traceability. The outputs summarize biomedical contexts and do not by themselves establish glycan functions or mechanisms.

The framework has three connected components. GlycoMeSH-BERT aligns glycan and MeSH representations in a shared space to infer associations beyond directly observed literature links (Fig. 1b). Literature-derived and predicted associations are integrated in GlycoMeSH-DB, expanding 27,563 associations between 3,061 glycans and 2,174 MeSH terms to 789,627 associations between 26,954 glycans and 20,302 terms (Fig. 1c). GlycoMeSH-EA uses this database for glycan-set enrichment with source PMIDs, prediction scores and user-defined score filtering (Fig. 1d).

The framework is intended to organize biomedical contexts associated with glycans, not to assign validated molecular functions. The three components are separable in use: GlycoMeSH-DB can be queried without running the inference model, and the enrichment workflow operates on any list of registered glycan structures. We therefore evaluated separately whether the model recovered held-out literature links, broadened the candidate MeSH space and retained motif-level semantic coherence before applying the resulting resource to glycan-set analysis.

Literature-derived MeSH terms provide informative but incomplete glycan contexts We constructed literature-derived glycan–MeSH annotations by linking GlyGen glycan– publication records to the MeSH terms assigned to each publication and filtering geographical, educational and highly prevalent non-specific terms (Methods). Because MeSH is assigned at the publication level, each pair records a biomedical context in which a glycan was reported; it does not by itself establish a direct mechanistic glycan–function relationship.

For example, G18982QA was reported in nigrostriatal tissue from patients with Parkinson’s disease^22^ and was linked to Corpus Striatum, Parkinson Disease, Substantia Nigra, Neurodegenerative Diseases and Lewy Body Disease (Supplementary Fig. 1a). G51367TM was reported across studies of hepatocyte alkaline-phosphatase glycosylation and pregnancy-related or trophoblastic conditions^23–28^, yielding distinct disease, phenotype, molecular and biological-context terms (Supplementary Fig. 1b and Supplementary Table 1).

Coverage was nevertheless sparse and heterogeneous. Of 2,174 curated MeSH terms, 27.5% were linked to one glycan and 74.0% to no more than ten glycans (Supplementary Fig. 1c). The 20 most frequent terms also included broad or experimental contexts, including Protein Structure, Tertiary, Hydrolysis, Trypsin, Ligands and Milk, Human (Supplementary Fig. 1d). Thus, literature-derived MeSH terms contain useful context but do not provide sufficient glycan coverage for general enrichment analysis.

### Inferring glycan–MeSH associations with GlycoMeSH-BERT

GlycoMeSH-BERT aligns pretrained glycan and MeSH embeddings by cross-modal contrastive learning. GlycanGT encoded branched glycan structures^13^, and four biomedical language models were evaluated for MeSH representation^14–17^, with SapBERT selected on performance (Supplementary Fig. 2a). Cosine similarity in the aligned space was used to rank candidate MeSH terms for each glycan (Fig. 2a).

**Fig. 2:**
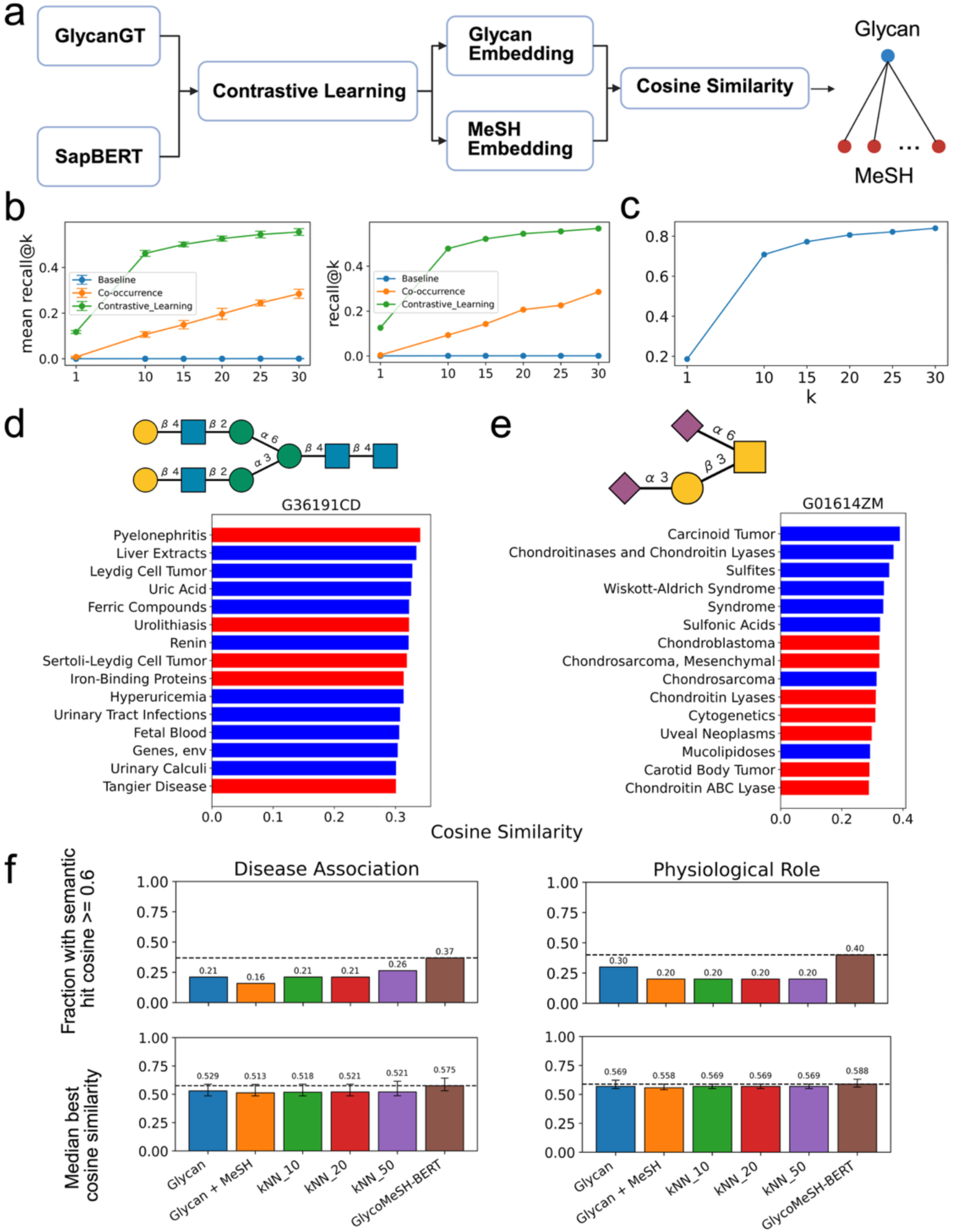
GlycoMeSH-BERT retrieves literature-derived associations and provides open-vocabulary motif-level semantic coverage. a, Overview of GlycoMeSH-BERT. Glycan and MeSH embeddings are aligned in a shared representation space using contrastive learning, and glycan–MeSH associations are ranked by cosine similarity. b, Recall@k comparison between baseline models and GlycoMeSH-BERT. Left, mean ± s.d. across five-fold cross-validation folds (n = 5). Right, held-out glycan test set. c, Recall@k in the held-out glycan test set after excluding glycans with recall@10 = 0. d,e, Top 15 MeSH terms for G36191CD (d) and G01614ZM (e). Blue denotes terms present in the current literature-derived document-level annotations; red denotes high-ranked terms absent from that curation and therefore not established as true positives. f, Motif-level semantic evaluation using manually curated motif-associated physiological and disease descriptions from Essentials of Glycobiology. The descriptions and predicted MeSH terms were compared in SapBERT space (semantic hit, best cosine similarity ≥ 0.60; error bars, interquartile ranges). This evaluation is independent of the training labels but uses the same text-embedding space as training and is therefore not fully model-independent; candidate-count–controlled comparisons are provided in Supplementary Table 11.

In glycan-level train–test splits, GlycoMeSH-BERT reached ∼50% recall@10 and ∼60% recall@30 for held-out literature-derived associations, outperforming direct cosine-similarity and co-occurrence baselines (Fig. 2b). Precision, F1 score, mean reciprocal rank and hit rate showed the same advantage over these non-aligned baselines (Supplementary Fig. 2b,c and Supplementary Tables 2 and 3). This experiment tests recovery of known document-level links, not the accuracy of novel predicted associations.

### Generalization and label incompleteness in GlycoMeSH-BERT predictions

Performance was comparable when test glycans were stratified by structural similarity to the training set, with only a small decrease in the low-similarity group (Supplementary Fig. 3 and Supplementary Table 4). This pattern indicates that held-out recovery was not explained solely by transfer from the most structurally similar training glycans.

Glycans with recall@10 = 0 had lower prediction scores than those with recall@10 > 0 (two-sided Mann–Whitney U test, P = 6.0 × 10−73 for the top 50 and P = 2.3 × 10−68 for the top 10) and fewer associated MeSH labels (P < 1 × 10−13; Supplementary Fig. 4 and Supplementary Table 5). Excluding these cases increased recall to ∼70% at 10 and ∼80% at 30 (Fig. 2c).

Zero recall can therefore coincide with both low model confidence and limited reference labels. Because missing literature labels are not verified negatives, these analysescannot determine how many unlabelled predictions are correct; they instead define label incompleteness as a constraint on benchmark interpretation.

### Representative predictions illustrate plausible unlabelled contexts

For G36191CD, GlycoMeSH-BERT recovered labelled terms including Liver Extracts, Leydig Cell Tumor, Uric Acid and Renin, while ranking the unlabelled terms Pyelonephritis and Urolithiasis highly (Fig. 2d). The glycan has been reported to bind Renin^29^, which is associated with pyelonephritis and urolithiasis^30,31^, providing a literature-based route by which these terms could arise.

For G01614ZM, the model recovered Carcinoid Tumor, Chondroitinases and Chondroitin Lyases, Sulfites and Wiskott–Aldrich Syndrome, and additionally ranked Chondrosarcoma and Chondroitin (Fig. 2e). Previous work linked chondroitin-binding glycans to chondrosarcoma^32^, making these unlabelled terms plausible but not independently validated associations.

These examples show that some absent labels have literature-consistent interpretations and motivate GlycoMeSH-BERT as an annotation-expansion model rather than a closed-label classifier. They do not estimate the overall precision of unlabelled predictions, which requires systematic literature review or experimental validation.

### Open-vocabulary inference expands MeSH coverage while retaining motif-level semantic agreement

We compared GlycoMeSH-BERT with two supervised multilabel MLP models and k-nearest-neighbour (kNN) label transfer using the same glycan-level split and ranking metrics (Supplementary Fig. 5a). The MLP and kNN models predict labels observed during training, whereas GlycoMeSH-BERT can score the full MeSH vocabulary; the comparison therefore separates closed-label recovery from open-vocabulary coverage.

The non-contrastive models matched or exceeded GlycoMeSH-BERT on literature-label recall: both MLPs exceeded 60% recall@10 and 80% recall@30, and kNN reached ∼60% and ∼70%, respectively (Supplementary Fig. 5b and Supplementary Tables 6 and 7). Thus, contrastive learning did not improve this retrieval metric. Coverage differed sharply: among aggregated top-100 predictions, GlycoMeSH-BERT generated 28,309 unique terms, compared with 6,285 for the glycan-only MLP, 1,568 for the glycan+MeSH MLP and 1,494– 1,709 for kNN (Supplementary Fig. 5c,d and Supplementary Tables 8 and 9).

The same pattern was evident within MeSH hierarchies. The glycan-only MLP retained broad parent-category coverage but recovered 947 Disease terms, compared with 4,073 for GlycoMeSH-BERT, with lower term diversity across neurological and immune-related categories (Supplementary Fig. 5e,f and Supplementary Table 9). This analysis quantifies breadth within biomedical categories; it does not establish the correctness of each additional term.

To test whether broader coverage retained glycan-related meaning, we compared predicted terms with 29 motif–description records manually curated from Essentials of Glycobiology, independently of the literature-derived training labels (Supplementary Table 10). The retained benchmark comprised 10 physiological-role and 19 disease-association records. At matched candidate counts, GlycoMeSH-BERT and the non-contrastive baselines showed comparable best semantic agreement. An apparent advantage emerged only when more candidate terms were considered and scaled with candidate number (Fig. 2f and Supplementary Table 11). Agreement was measured with SapBERT cosine similarity, the same text-embedding space used during training, and is therefore not a model-independent measure of semantic accuracy.

Together, these results define a trade-off relevant to resource construction. Closed-label models efficiently recover dominant known annotations, whereas GlycoMeSH-BERT broadens the MeSH space while retaining comparable motif-level semantic agreement at matched candidate counts. This evidence supports its use as an open-vocabulary coverage engine for GlycoMeSH-DB.

### Construction of GlycoMeSH-DB

We retrained GlycoMeSH-BERT on the full annotation dataset and scored GlyTouCan glycans without ambiguous notation. A cosine-similarity threshold of 0.3 was selected from sensitivity analyses of association counts, retention of literature-derived pairs and the ratio of unmatched to matched predictions (Supplementary Fig. 6a,b). After integration with literature-derived annotations and deduplication, GlycoMeSH-DB contained 789,627 associations linking 26,954 glycans to 20,302 MeSH terms (Fig. 3a and Supplementary Table 12).

**Fig. 3:**
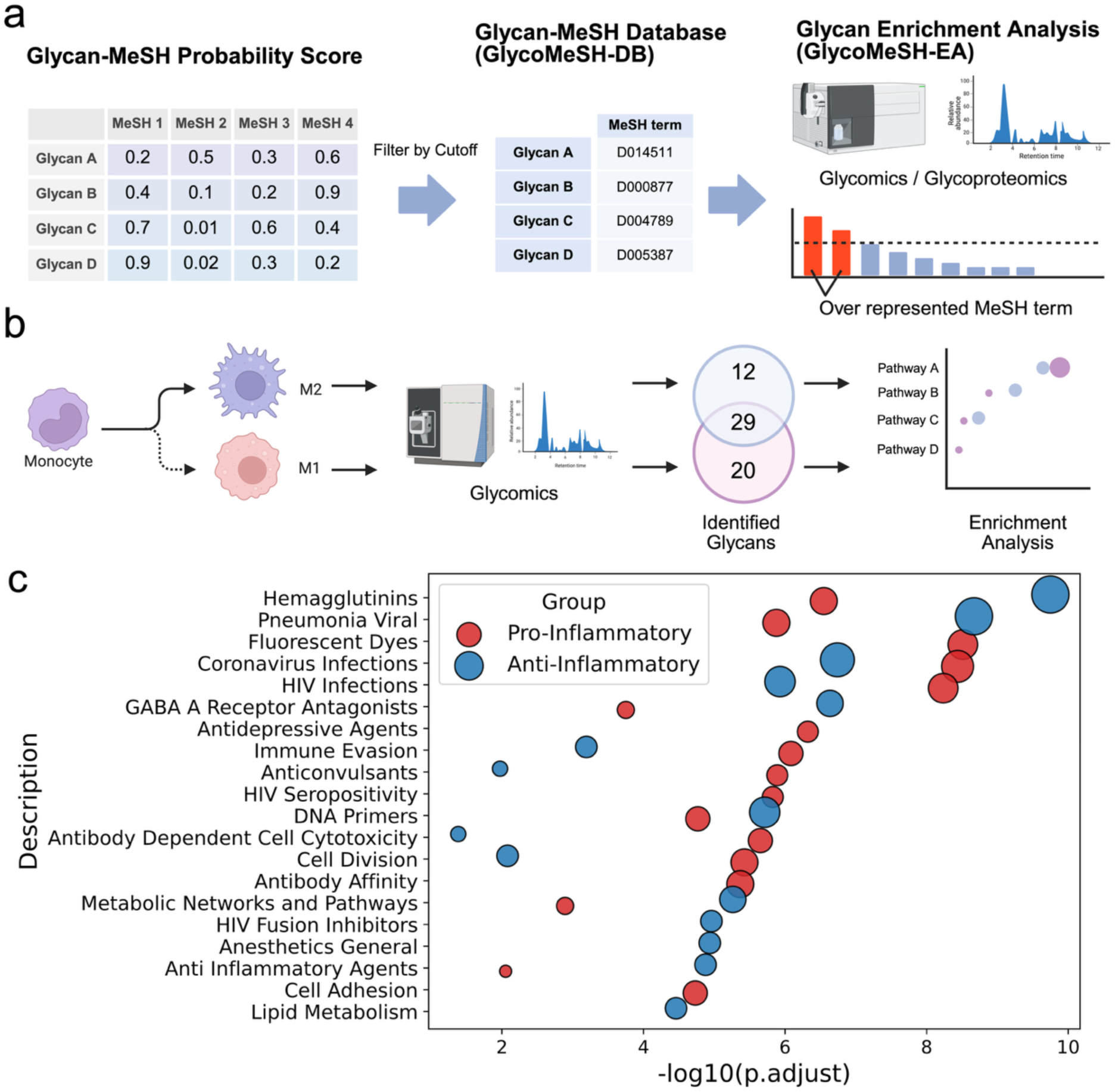
Construction of GlycoMeSH-DB and illustrative application of GlycoMeSH-EA to human glycomics data. a, Construction of GlycoMeSH-DB by thresholding the GlycoMeSH-BERT cosine-similarity matrix and integrating predicted associations with literature-derived annotations. At the 0.3 operational threshold, the database represents annotation coverage rather than an individually validated knowledge set. b, Illustrative GlycoMeSH-EA workflow for human macrophage glycomics. Glycans detected in the pro-inflammatory and anti-inflammatory states were used as query sets. c, Enrichment results for the pre-selected display categories. ORA used the hypergeometric test (enricher, clusterProfiler), all glycans in the analysis as background and Benjamini–Hochberg FDR control across the full MeSH vocabulary. The categories shown were selected in advance for display only, and unrestricted results are provided in Supplementary Table 14; these results demonstrate use of the method and are not an independent validation of biological specificity.

Literature PMIDs and prediction cosine scores are retained for every association, so that users can distinguish evidence sources and apply stricter score filters. The threshold itself is operational rather than a calibrated probability of correctness: 789,627 denotes annotation coverage at 0.3, and individual predicted associations were not separately validated for precision. Because score calibration has not been established, filtering can rank evidence but does not convert a cosine score into a probability of correctness.

### Illustrative application to human macrophage glycomics

We applied GlycoMeSH-EA to a human glycomics dataset whose PMID was absent from the literature-derived annotation set. CD14+ monocytes were differentiated into pro-inflammatory M1 and anti-inflammatory M2 macrophages^33^, and glycans were identified with CandyCrunch^9^ (Supplementary Table 13). M1 glycans were enriched for infection-and immune-response-related terms, whereas M2 glycans included Anti-Inflammatory Agents (Fig. 3b,c and Supplementary Table 14). Literature-derived annotations covered only one detected glycan (G65890UA), precluding comparable enrichment with literature links alone. Enrichment was tested across the full MeSH vocabulary, with Benjamini–Hochberg FDR control applied over all tested terms, and the complete unrestricted results are provided in Supplementary Table 14. Fig. 3c displays the pre-selected parent categories Immune System Phenomena, Cell Physiological Phenomena, Infections, Metabolism, and Chemical Actions and Uses. Because these display categories were chosen for biological relevance, the figure illustrates workflow output rather than an independent validation of biological specificity.

### Illustrative application to Alzheimer’s disease glycoproteomics

We next analyzed human and mouse Alzheimer’s disease (AD) glycoproteomics datasets whose PMIDs were absent from the literature-derived annotation set. This independence concerns overlap with the training literature; it does not constitute independent validation of predicted biology.

In N-linked glycoproteomics data from postmortem brains of patients with AD and cognitively normal individuals^34^, Glyco-Decipher identified glycopeptides^9^ (Supplementary Table 15).

Enrichment was again tested across the full MeSH vocabulary. Among the pre-selected Nervous System Diseases, Neurodegenerative Diseases and Mental Disorders categories displayed in Fig. 4a, the AD group showed Cognition Disorders, Dementia and Neuroinflammatory Disease, whereas the control group included less dementia-specific terms such as Adie Syndrome; results for all tested terms are provided in Supplementary Table 16.

**Fig. 4:**
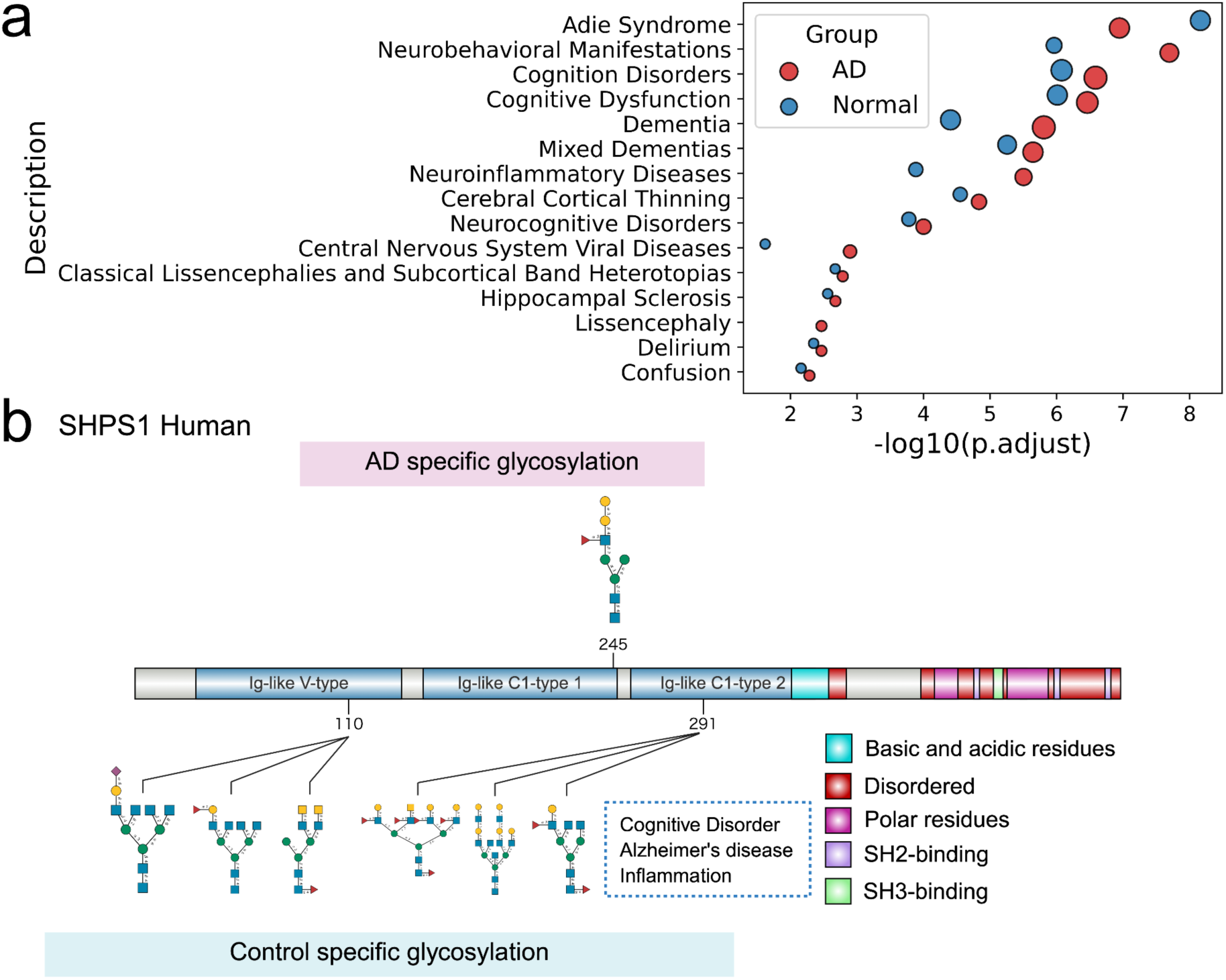
Illustrative application of GlycoMeSH-EA to human Alzheimer’s disease glycoproteomics data. a, GlycoMeSH-EA enrichment results for glycans detected in Alzheimer’s disease and cognitively normal control samples. ORA used the hypergeometric test (enricher, clusterProfiler), all glycans in the corresponding analysis as background and Benjamini– Hochberg FDR control across the full MeSH vocabulary. Displayed nervous-system categories were selected in advance for display only, and unrestricted results are provided in Supplementary Table 16, so the panel is illustrative rather than an independent validation of disease specificity. b, Descriptive domain-level glycosylation patterns of SHPS1. SHPS1 was selected post hoc as a hypothesis-generating example; the control-specific glycans shown in the Ig-like V-type domain were not tested for domain-localization specificity and do not establish a mechanism.

In hippocampal glycoproteomics from APP/PS1 and control mice^35^, Cognitive Dysfunction and Dementia were enriched in both groups, whereas Cerebral Cortical Thinning, Delirium, Confusion and Memory Disorder were enriched in the AD model (Supplementary Fig. 7a and Supplementary Tables 17 and 18). These patterns are consistent with reported hippocampal atrophy and AD-like behavioural abnormalities in APP/PS1 mice^36^, but consistency with prior knowledge is not an independent specificity test.

Across the full MeSH vocabulary, ORA using a hypergeometric test with Benjamini– Hochberg FDR control identified 177–180 significant MeSH terms across the human groups and 128–130 across the mouse groups, distributed across many MeSH parent categories beyond those displayed (Supplementary Tables 16 and 18).

Using literature-derived glycan–MeSH links alone did not recover AD- or dementia-related neurological terms in either dataset (Supplementary Tables 16 and 18). Protein-side comparisons yielded too few significantly altered glycoproteins for robust gene-centric enrichment (Supplementary Table 19). These underpowered comparators show the additional contexts accessible from the expanded glycan annotation layer; they do not validate the glycan-side enrichments and are best regarded as orthogonal analyses.

As a post hoc, hypothesis-generating example, we examined SHPS1/SIRPα, an AD-associated membrane glycoprotein encoded by SIRPA^37^. Its Ig-like V-type domain mediates CD47 binding^38^. In both datasets, control-specific glycans were observed in this domain descriptively, without a statistical test of domain-localization specificity, and co-occurred with terms including Cognitive Aging and Neuroinflammatory Diseases (Fig. 4b, Supplementary Fig. 7b and Supplementary Tables 20 and 21). Prior evidence that SIRPα glycosylation modulates CD47 binding motivates—but does not test—the hypothesis that altered domain glycosylation could influence CD47–SIRPα signalling in AD.

The macrophage and AD analyses therefore illustrate how GlycoMeSH-EA surfaces glycan-centric biomedical contexts at dataset and glycoprotein levels, including contexts that the literature-derived layer alone did not recover. Enrichment testing and FDR control were unrestricted, and complete results are provided in the Supplementary Information; the pre-selected categories affect only which terms are displayed. Because those display categories were chosen for biological relevance and no specificity negative control was analyzed, these applications demonstrate workflow use, not general biological validity.

### GlycoMeSH-EA: a web-based glycan-set enrichment tool

GlycoMeSH-EA implements GlycoMeSH-DB as a web interface for glycan-set enrichment (Fig. 5). Users submit GlyTouCan IDs, examine enriched terms through MeSH parent categories, trace literature-derived associations to source PMIDs and inspect cosine scores for predicted associations. Score filtering lets users trade coverage for confidence while keeping literature and model-derived evidence distinguishable. The interface therefore exposes both enriched terms and the association-level provenance contributing to them, allowing outputs to be audited rather than treated as opaque model predictions. Because the query unit is a list of GlyTouCan accessions, any glycomics or glycoproteomics workflow that reports registered structures can be analyzed without first collapsing the structures into motif or composition categories, and without regard to the identification software used upstream. The datasets analyzed here were processed by two different identification tools, and both structure lists were submitted through the same interface without reformatting.

**Fig. 5:**
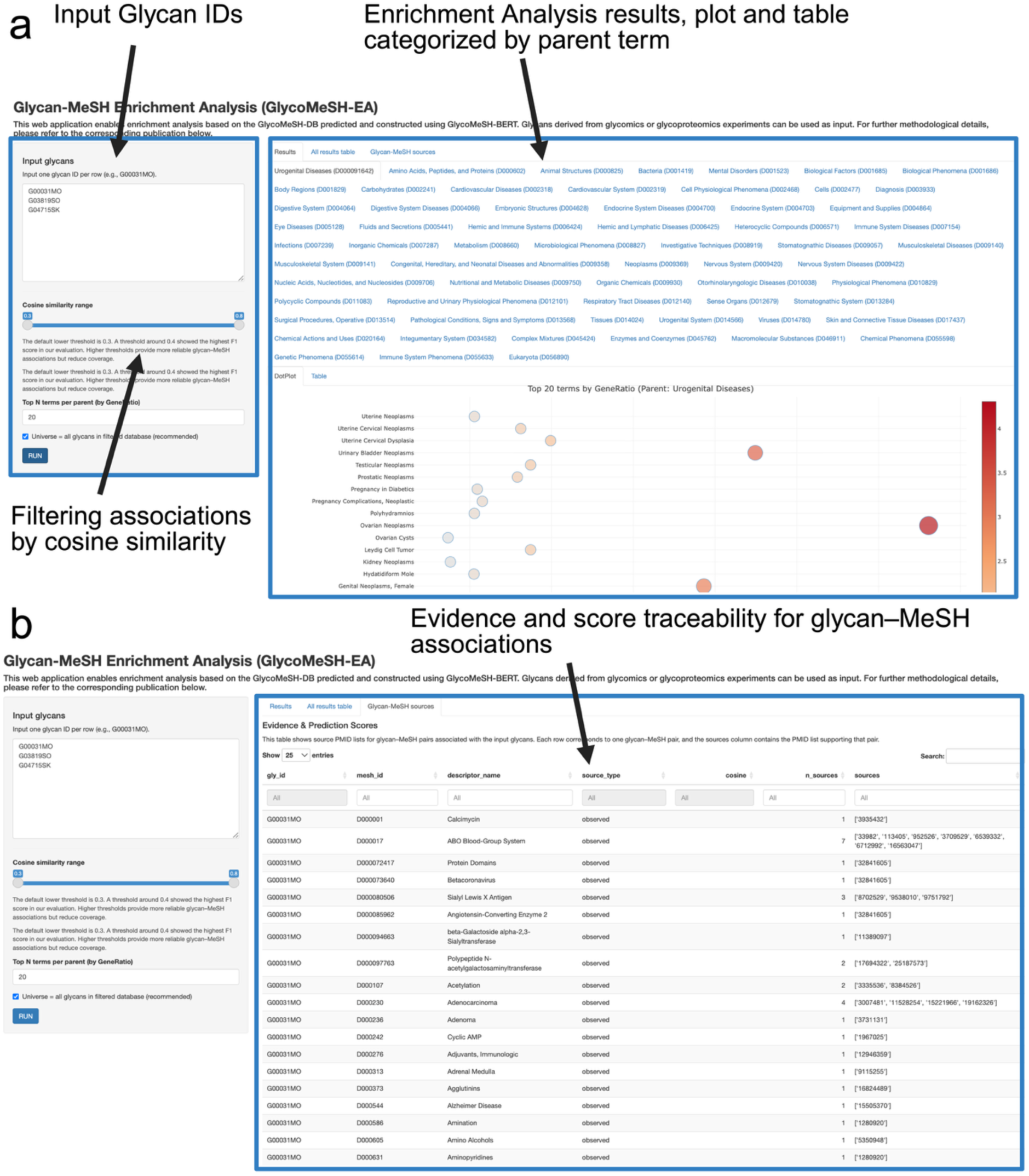
Web-based implementation of GlycoMeSH-EA. **a**, GlycoMeSH-EA interface for glycan-centric enrichment analysis using GlycoMeSH-DB. Users can input a set of glycans (e.g., GlyTouCan IDs) to identify enriched MeSH terms and explore results organized by hierarchical MeSH categories. **b**, Evidence and score traceability panel. The table enables users to inspect source PMIDs for literature-derived glycan–MeSH annotations and cosine similarity scores for prediction-derived associations.

## Discussion

GlycoMeSH provides the missing annotation layer between glycan structure and reusable biomedical context. It integrates open-vocabulary inference, a traceable glycan–MeSH database and glycan-set enrichment so that detected glycans can be analyzed directly rather than only through structural summaries or gene and protein proxies. At the association level, source PMIDs and prediction scores preserve the distinction between observed literature evidence and model inference. This traceability is central to reuse because users can audit the evidence supporting an enriched term and adjust prediction-score thresholds. Rather than assigning validated molecular mechanisms, the contribution is a resource and analysis framework that makes glycan-associated contexts queryable and auditable.

The model comparisons expose the central trade-off. Closed-label MLP and kNN models recovered sparse literature labels as well as or better than GlycoMeSH-BERT, but produced a narrower vocabulary. GlycoMeSH-BERT broadened MeSH coverage and retained comparable motif-level semantic agreement when candidate counts were matched. For a resource intended to support enrichment, breadth matters because an annotation space dominated by a small set of frequent labels limits the contexts that can be queried.

Breadth is not sufficient by itself, however, and should be coupled to confidence filtering and evidence inspection. The results therefore support GlycoMeSH-BERT as a coverage engine under incomplete labels. The motif analysis is independent of the training labels but shares the text-embedding space used during training, and therefore measures semantic agreement rather than accuracy.

GlycoMeSH complements, rather than replaces, structural databases, glycan ontologies, glycoenzyme pathways, host–pathogen resources and lectin classifications^41–46^. It also differs from enzyme-mediated transfer of Gene Ontology terms, such as GlycoEnzOnto^44^, because its query unit is the glycan structure itself. This distinction matters when a signal is carried by site-specific glycoforms or when too few proteins change for gene-centric enrichment. A controlled glycan-side vocabulary also permits the same structure-level query to be reused across experiments, datasets and laboratories without redefining ad hoc motif categories for each study. This lowers the barrier for groups that do not develop annotation methods themselves: a laboratory reporting a glycan list can obtain biomedical-context terms, inspect the evidence behind each one and set its own score threshold without retraining or reimplementing any component. The macrophage and AD applications show how the interface can expose such contexts, but their pre-selected categories and limited datasets make them demonstrations of utility rather than validation of general biological specificity.

Several boundaries define current use. Literature-derived pairs are incomplete, heterogeneous and reporting-biased, and missing labels are not true negatives. Glycan-identification uncertainty, including unresolved linkage, branching and positional isomers, propagates to annotation and enrichment. MeSH also retains broad technical or chemical contexts, and GlycoMeSH-DB’s 789,627 associations represent coverage at an operational 0.3 threshold rather than individually validated knowledge. ORA assumes independent query glycans and may be anti-conservative when related glycans co-occur and share annotations; application results should therefore be interpreted descriptively. The application datasets cover only immune and neurodegenerative settings, and the usability of the web interface has not been formally evaluated. Future work should calibrate prediction confidence, quantify precision across score ranges, use degree-preserving null models, add specificity negative controls across more biomedical domains and formally evaluate the web interface. Such validation would convert coverage into calibrated evidence without changing the resource’s core separation of literature and inferred associations. Within the present boundaries, GlycoMeSH makes glycan-set interpretation traceable, reusable and directly accessible.

## Methods

### Data sources

Glycan data were obtained from GlyCosmos^47^. We collected GlyTouCan IDs, IUPAC-condensed representations, and the corresponding PMID information for glycans available as of January 30, 2026. MeSH is a controlled vocabulary maintained by the National Library of Medicine (NLM) for indexing biomedical and life science literature. Each publication is assigned multiple MeSH terms through automated mapping and curation. MeSH descriptor data were obtained from the 2026 release distributed by the NLM.

### Data processing

For the glycan dataset, entries lacking IUPAC-condensed representations were excluded, resulting in 69,554 glycans for analysis. To obtain feature representations for each glycan structure, embeddings were computed using GlycanGT^13^. GlycanGT is a glycan structure representation learning model based on a graph transformer architecture^48^, which takes an IUPAC-condensed representation as input and outputs a 768-dimensional glycan embedding. GlycanGT weights were also kept fixed, and only the downstream projection heads were optimized in this study. For MeSH terms, we generated two types of text input: one consisting of the descriptor name alone, and the other consisting of the descriptor name, synonyms, and definition concatenated into a single label text (Supplementary Table 22). The former was input to SapBERT^17^, whereas the latter was input to BioBERT^14^, PubMedBERT^15^, and MedCPT^16^. For SapBERT, BioBERT, and MedCPT, the embedding of each MeSH term was obtained from the [CLS] token representation. For PubMedBERT, mean pooling over token embeddings was used. All language model weights were kept fixed during downstream training. For all models, a 768-dimensional embedding representation was obtained for each MeSH term. SapBERT is a pretrained model specialized for biomedical entity representation learning, whereas BioBERT, PubMedBERT, and MedCPT are language models pretrained on biomedical text corpora such as PubMed. Glycan–MeSH annotations were constructed by integrating glycan–PMID associations obtained from GlyGen^49^ with the MeSH terms assigned to each PMID. Highly general MeSH terms that were assigned uniformly across many glycans were removed, because such terms could artificially inflate prediction performance while contributing little to the functional interpretation of glycans. Specifically, terms associated with more than 200 glycans were regarded as non-specific and excluded. This cutoff was chosen empirically to remove highly prevalent descriptors with limited discriminative value for glycan-level functional interpretation. Furthermore, terms not relevant to the aim of this study, such as those related to education or geographical information, were manually curated and removed. These literature-supported glycan–MeSH associations served as a broad-sense positive control set for model evaluation: each association is supported by published evidence at the document level, even though it does not necessarily reflect a mechanistically validated glycan–function relationship. After these filtering procedures, 3,061 glycans were linked to 2,174 MeSH terms, yielding a total of 27,563 glycan–MeSH annotations (Supplementary Table 1).

### GlycoMeSH-BERT architecture

In this study, we developed a cross-modal joint embedding model to align glycan embeddings and MeSH embeddings in a shared latent space. The inputs consisted of precomputed glycan embedding vectors and MeSH embedding vectors. A separate projection head was applied to each modality; each projection head comprised a multilayer perceptron with a linear layer, a ReLU activation, dropout, and a second linear layer,

followed by L2 normalization. Let *X*_g_ ∈ *R^dg^* and *X_m_* ∈ *R^dm^* denote the input embeddings of glycan *g* and MeSH term *m*, respectively. The projected representations were defined as follows:

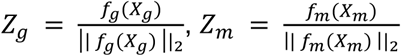

where *f*_g_ and *f_m_* denote the projection heads for the glycan and MeSH modalities, respectively. To account for the fact that a single glycan can be associated with multiple MeSH descriptors, we adopted a multi-positive contrastive objective based on the StableRep-style formulation^11^. In each mini-batch, multiple positive MeSH terms were sampled for each glycan anchor. For a given glycan anchor *i*, similarity to all candidate MeSH embeddings within the batch was computed using temperature-scaled cosine similarity as follows:

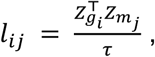

where *τ* denotes the temperature parameter. The target distribution *p_i_*_j_ was defined as a uniform distribution over the positive MeSH set associated with anchor *i*, and 0 otherwise. Training was performed by minimizing the cross-entropy between this target distribution and the predicted distribution obtained by applying softmax to the similarity scores:

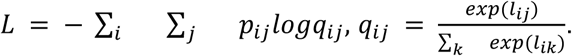

This formulation encouraged each glycan representation to be positioned close to all corresponding MeSH representations in the shared latent space.

### Model training and hyperparameter optimization

Model training was performed using a two-stage optimization strategy (coarse-to-fine training). In the first stage (stage 1), a relatively large learning rate was used to explore the parameter space broadly. In the second stage (stage 2), the model parameters obtained from stage 1 were used as initialization, and a smaller learning rate was applied for local refinement. Optimization was performed using AdamW with weight decay to suppress overfitting. Gradient clipping was additionally applied to stabilize training. Mini-batches were constructed using a multi-positive scheme in which glycans were sampled first, and multiple positive MeSH terms were assigned to each glycan. Thus, each batch consisted of multiple glycan anchors, with each anchor paired with multiple positive MeSH terms. Pairs derived from the same glycan were treated as belonging to the same group, and the positive set was defined accordingly within the loss function.

Model performance was evaluated using retrieval metrics on glycan-level data splits, including hit@k, recall@k, precision@k, F1 score, and mean reciprocal rank (MRR). These metrics were used to evaluate the model as a ranking task, in which MeSH terms were ranked for each glycan based on cosine similarity. Recall@k measures the fraction of true associated MeSH terms retrieved within the top-k predictions, precision@k evaluates the proportion of correct predictions among the top-k results, F1 score represents the harmonic mean of precision and recall, hit@k indicates whether at least one correct MeSH term is included in the top-k, and mean reciprocal rank (MRR) reflects the rank of the first correct prediction. These metrics are well-suited for multi-label retrieval settings, where each glycan may be associated with multiple MeSH terms. Hyperparameter search and model selection were both performed based on glycan-level splits, such that the same glycan was never included in both the training and evaluation datasets.

We adopted glycan-level rather than PMID-level splits because the primary objective was to evaluate recovery of MeSH associations for glycans not observed during training. Multiple glycans can be reported in one publication and share MeSH descriptors, reflecting the structure of the literature-derived annotations rather than simple duplication. PMIDs were therefore allowed to overlap between training and test sets, with an overlap rate of approximately 50%. PMID identifiers and publication-level information were not model inputs; the model received only glycan-structure and MeSH-term embeddings. PMID overlap therefore does not constitute input-level leakage of held-out glycan structures or their labels. It does not, however, exclude label-side dependence: related glycans co-reported in one publication can inherit correlated MeSH labels. A PMID-disjoint sensitivity analysis would more directly quantify this residual dependence and remains future work.

Hyperparameter optimization followed a two-step strategy consisting of random search and Bayesian optimization with Optuna. First, for each of the four MeSH embedding types (BioBERT, MedCPT, PubMedBERT, and SapBERT), hyperparameters including learning rate, temperature parameter, dropout, weight decay, batch configuration (batch size and number of positives), and projection head dimensions were sampled from log-uniform or uniform distributions, and broad exploration was performed by random search. Next, Optuna-based hyperparameter optimization was conducted for the best-performing embedding setting, namely SapBERT (Supplementary Fig. 2a). For this step, the dataset was split at the glycan level into a fixed test set (15% of the data) and a development set (the remaining 85%), and 5-fold cross-validation was performed on the development set. For each trial, the mean cross-validation performance (recall@25) was maximized as the objective function, and pruning based on intermediate results was used to improve search efficiency. After the optimal hyperparameters were identified, a model was retrained on the entire development set without using a validation set, and its performance was evaluated on the fixed test set.

For comparison, two non-contrastive baselines were evaluated under the same train/test split and retrieval metrics. In the baseline model, glycan–MeSH relationships were estimated directly from cosine similarity in the original embedding spaces of glycans and MeSH terms. In addition, we evaluated a co-occurrence baseline, in which MeSH terms were ranked solely according to their frequency in the training annotations, independent of glycan structure.

After model selection and benchmark evaluation, the final model used for GlycoMeSH-DB construction was trained separately on the full glycan–MeSH annotation dataset using the selected hyperparameters. In this full-data training procedure, all available pairs were used for optimization, and no validation split, early stopping, or model selection based on validation performance was applied. The final checkpoint obtained after stage 2 was used for downstream inference and GlycoMeSH-DB construction. Final hyperparameters and reproducibility settings for GlycoMeSH-BERT and the MLP-based models are summarized in Supplementary Table 23.

### Analysis of model generalization and annotation sparsity

To assess model generalization, test glycans were stratified according to their structural similarity to glycans in the training set. Specifically, cosine similarity was calculated between the GlycanGT embedding of each test glycan and those of all glycans in the training set. For each test glycan, the 75th percentile of the resulting similarity distribution was used as a summary score of structural relatedness to the training set. Test glycans were then grouped into high- and low-similarity categories using a threshold of 0.5 on this 75th- percentile similarity score, yielding 269 high-similarity and 173 low-similarity glycans. Prediction performance was then compared between the resulting groups. In addition, glycans with recall@10 = 0 were compared with those with recall@10 > 0 in the test set.

To evaluate differences in model confidence, the distribution of cosine similarity scores assigned to predicted glycan–MeSH pairs was compared between the two groups for the top 10 and top 50 ranked predictions using a two-sided Mann–Whitney U test. We also examined the relationship between prediction performance and annotation coverage by comparing the number of MeSH labels assigned to each glycan between the two groups using a two-sided Mann–Whitney U test. For qualitative assessment of model predictions, representative glycans were randomly selected from the test set, and their top-ranked MeSH terms were examined for agreement with known annotations and consistency with prior literature.

### Comparison with non-contrastive baseline models

Held-out recall against the curated literature-supported glycan–MeSH set (defined above as a broad-sense positive control) was used as the primary criterion for comparing GlycoMeSH-BERT against alternative non-contrastive approaches. To assess the methodological advantage of contrastive learning, we compared GlycoMeSH-BERT with non-contrastive baseline models, including two supervised multi-label MLP-based models and a non-parametric k-nearest neighbor (kNN) label-transfer baseline. The first model used only GlycanGT-derived glycan embeddings as input (glycan-only MLP), whereas the second used both glycan embeddings and SapBERT-derived MeSH embeddings (glycan+MeSH dual MLP). In both cases, prediction was formulated as multi-label classification over the full candidate MeSH space, and MeSH terms were ranked for each glycan according to the model output scores. For the glycan-only model, GlycanGT embeddings were input to a multilayer perceptron classifier that directly outputs logits for all candidate MeSH terms. For the glycan+MeSH model, separate MLP projection heads were applied to glycan and MeSH embeddings, and the score for each glycan–MeSH pair was computed as a scaled dot product in the projected latent space, which was used as the logit for multi-label classification over the full MeSH vocabulary. Projected embeddings were L2-normalized, and supervised multi-label learning was performed using BCEWithLogitsLoss over the full candidate MeSH space. Both MLP-based models used the same hyperparameter optimization and two-stage training procedure. Hyperparameter optimization was performed using a fixed glycan-level test split and 5-fold cross-validation on the development set, followed by Bayesian optimization with Optuna. The search space included hidden dimensions, dropout rate, learning rates for two-stage training, weight decay, batch size, positive-class weighting, and, for the dual-MLP model, projection dimension and modality-specific hidden dimensions. After hyperparameter selection, each model was retrained on the full development set without a validation split and evaluated on the fixed test set.

Training used a two-stage optimization scheme with AdamW and BCEWithLogitsLoss, with optional class balancing by positive weights. For fair comparison, both MLP-based models were trained and evaluated using the same glycan-level data partitioning strategy, the same fixed test split, and the same full candidate MeSH vocabulary as used for GlycoMeSH-BERT.

For the kNN label-transfer baseline, predictions were generated in the GlycanGT embedding space without model training. Glycan embeddings were L2-normalized, and cosine similarity was calculated between each query glycan and glycans in the training set.

For each query glycan, the top-k nearest training glycans were selected, and MeSH terms assigned to these neighbors were transferred to the query glycan. Each candidate MeSH term was scored by summing the cosine similarities of neighboring glycans annotated with that term. We evaluated k = 10, 20, and 50, and MeSH terms were ranked according to the resulting label-transfer scores. For the MLP-based models, MeSH terms were ranked according to output logits, which are monotonic with predicted probabilities. For the kNN baseline, MeSH terms were ranked according to label-transfer scores. All models were evaluated using the same ranking-based metrics, including hit@k, recall@k, precision@k, F1 score, and mean reciprocal rank (MRR). To assess the diversity of predicted functions, we quantified the number of unique MeSH terms appearing within the aggregated top 100 predictions across all glycans for each model. We also counted the frequency of occurrence of individual MeSH terms within these top-ranked predictions to evaluate whether predictions were broadly distributed or concentrated on a limited subset of terms.

To further evaluate diversity at the MeSH hierarchy level, predicted MeSH terms were mapped to their parent terms using the MeSH descriptor hierarchy, and the number of unique parent terms represented by each model was counted. In addition, to assess term-level diversity within disease-related categories, predicted MeSH terms assigned to Disease-related parent terms were grouped by parent category, and the number of unique terms within each parent category was compared between the GlycoMeSH-BERT and the glycan-only MLP model.

To evaluate semantic coherence with motif-associated biomedical descriptions beyond the literature-derived glycan–MeSH annotations, we used textbook-curated glycan motif annotations from Essentials of Glycobiology (Supplementary Table 10). We curated associations between IUPAC-condensed glycan motifs and textual descriptions of physiological roles or disease associations. Each description was embedded using SapBERT, with the same procedure used for MeSH descriptor names. For each retained motif–description record and each model, predicted MeSH terms were collected from glycans whose IUPAC-condensed structures contained the corresponding motif. Predictions were evaluated at rank cutoffs of 1, 5, 10, 20, 30, 50 and 100. For each cutoff, cosine similarity was calculated between the SapBERT embedding of the curated description and that of each predicted MeSH term, and the highest value was retained as the best semantic match. Summary statistics comprised the mean and median best similarity, interquartile range and the fraction of records with a semantic hit, defined as best cosine similarity ≥ 0.60. We initially curated 42 records: 16 physiological-role and 26 disease-association records.

Motif-containing glycans were identified by matching IUPAC-condensed motif strings against the analyzed GlyCosmos/GlyTouCan set. Records without an explicit motif string or a structure-matched glycan were excluded, leaving 29 records (10 physiological-role and 19 disease-association records); 13 were excluded because no structure-matched glycan was found. GlycoMeSH-BERT and each baseline were compared across matched records at each cutoff using two-sided Wilcoxon signed-rank tests, separately for the two description types. P values were adjusted by the Benjamini–Hochberg procedure across description types, rank cutoffs and baseline comparisons. Because both curated descriptions and predicted MeSH terms were represented with SapBERT, this evaluation is independent of the literature-derived training labels but not of the text-embedding space used during training. Full motif-level results are provided in Supplementary Table 11.

### Construction of GlycoMeSH-DB and implementation of GlycoMeSH-EA

Using the final trained model, all glycans derived from GlyCosmos that lacked ambiguous symbols such as “/” or “?” were input to GlycoMeSH-BERT, and cosine similarity was calculated between glycan embeddings and MeSH embeddings. Predicted glycan– MeSH associations were then defined by thresholding the cosine similarity scores. To determine the threshold for GlycoMeSH-DB construction, we evaluated a series of cosine similarity cutoffs using glycans that were shared between the literature-derived annotation set and the full-model prediction output. For each threshold, predicted glycan–MeSH pairs were compared with known literature-derived pairs, and we quantified both the number of retained known associations and the number of unmatched predicted pairs. Here, unmatched pairs were defined as predicted glycan–MeSH pairs that were not present in the current literature-derived annotation set. We then examined the trade-off between retention of known associations and expansion of the glycan–MeSH annotation space, including the ratio of unmatched predictions to matched true-positive predictions (Unmatched/TP), the total number of predicted pairs, and the coverage of glycans with at least one assigned term. Based on this analysis, a threshold of 0.3 was selected because it preserved nearly all known associations while allowing substantial expansion of glycan–MeSH coverage without a disproportionate increase in unmatched predictions (Supplementary Fig. 6). Predicted glycan–MeSH associations passing this threshold were integrated with literature-derived annotations, and duplicate glycan–MeSH pairs were removed to construct the final GlycoMeSH-DB (Supplementary Table 12). In addition to glycans registered in GlyTouCan, glycan structures detected in the application datasets but not assigned GlyTouCan accessions were also embedded using GlycanGT, and their associated MeSH terms were inferred using GlycoMeSH-BERT for downstream analysis.

To facilitate broader use of GlycoMeSH-DB, we implemented GlycoMeSH-EA as a web-based graphical user interface using R Shiny. The tool performs GlycoMeSH-DB-based enrichment analysis from GlyTouCan IDs with the enricher function in clusterProfiler and returns enriched MeSH terms associated with the input glycan set. Parent-term relations enable hierarchical exploration of enriched MeSH categories. For traceability, the interface reports the evidence source for each glycan–MeSH association. Literature-derived associations link to source PMIDs, whereas prediction-derived associations include GlycoMeSH-BERT cosine-similarity scores. Users can specify a score range before enrichment; the default lower threshold is 0.3, and higher thresholds prioritize score at the expense of coverage. The interface therefore preserves the distinction between literature-supported and prediction-derived associations without treating the prediction score as a calibrated probability.

### Application of GlycoMeSH-EA to glycomics and glycoproteomics datasets

GlycoMeSH-EA was applied to previously published glycomics and glycoproteomics datasets using GlycoMeSH-DB as TERM2GENE. The datasets comprised human macrophage glycomics^33^, postmortem human AD brain glycoproteomics^34^ and APP/PS1 mouse hippocampal glycoproteomics^35^. CandyCrunch^9^ was used for glycan identification and relative quantification, and Glyco-Decipher^8^ for glycopeptide identification and glycan assignment. Only CandyCrunch strong-evidence glycans were analyzed; internal identifiers were assigned where GlyTouCan accessions were unavailable. In the source macrophage study, CD14+ monocytes were differentiated with GM-CSF and polarized with IFN-γ/LPS or IL-4/IL-13. Glycans detected in the pro- and anti-inflammatory states were used as query sets, and displayed terms were restricted to Immune System Phenomena, Cell Physiological Phenomena, Infections, Metabolism, and Chemical Actions and Uses. The human AD data were from postmortem frontal cortex, and the mouse data from hippocampi of 9-month-old APP/PS1 and age-matched control mice. Displayed terms were restricted to Nervous System Diseases, Neurodegenerative Diseases, and Mental Disorders. These category restrictions were applied only to visualization, after enrichment testing and FDR control across the full MeSH vocabulary, and define the interpretive scope of the application results.

To assess the contribution of prediction-derived annotations, enrichment analyses were also performed using only the curated literature-derived glycan–MeSH annotations. For this comparison, the same query glycan sets, background glycans, enrichment function, display category filters, and multiple-testing correction procedure were used, but the TERM2GENE mapping was restricted to literature-derived glycan–MeSH associations.

For conventional protein-side interpretation, protein-level abundance matrices generated from the human and mouse glycoproteomics datasets were analyzed between disease and control groups using the same procedure. Proteins with zero variance across samples were removed. For each protein, the difference in mean log2 abundance between disease and control groups was calculated as log2 fold change, and differential abundance was assessed using a two-sided Welch’s t-test. P values were adjusted using the Benjamini–Hochberg method. Protein annotations were added using peptide-to-protein mapping information. Significantly altered proteins were then considered as candidates for conventional gene-centric enrichment analysis; when too few proteins passed the significance threshold, downstream gene-centric enrichment analysis was considered insufficiently powered and was not interpreted.

As an illustrative, hypothesis-generating example of glycoprotein-level interpretation, SHPS1 was selected post hoc because it is a membrane glycoprotein with disease relevance in Alzheimer’s disease and showed control-specific glycoforms in the analyzed datasets; this descriptive selection was not based on a pre-specified statistical test. Glycans assigned to SHPS1 were compared across conditions. Identified glycopeptides were mapped to the corresponding protein domains, including the Ig-like V-type domain, and interpreted in that structural context. Schematic illustrations of glycosylation sites and surrounding amino acid sequences on SHPS1 were prepared using IBS 2.0^50^.

### Statistical analyses

Statistical analyses were performed using Python and R. Glycomics and glycoproteomics enrichment analyses used the enricher function in clusterProfiler with GlycoMeSH-DB as TERM2GENE. Query sets comprised glycans detected in each condition, and the background universe comprised all glycans included in the corresponding analysis. ORA was performed across the full MeSH vocabulary represented in GlycoMeSH-DB. For each MeSH term, ORA used the hypergeometric distribution, with P values adjusted by the Benjamini–Hochberg procedure over all tested terms; category restrictions were applied only when selecting terms for display, after testing and correction. The hypergeometric test assumes independent sampling of query glycans. This assumption is not strictly met because structurally related glycans can co-occur and share MeSH labels; the test may therefore be anti-conservative. Enrichment results are consequently interpreted as descriptive and illustrative. Unless otherwise specified, tests were two-sided. Data visualization and machine-learning analyses used Python 3.12.11, PyTorch 2.5.1+cu121, Optuna 4.7.0, scikit-learn 1.7.0 and SciPy 1.15.3. Enrichment analyses used R 4.4.0 and clusterProfiler 4.14.6.

## Data availability

All data used in this study were obtained from publicly available sources. Glycan structures, GlyTouCan accessions, IUPAC-condensed representations and the associated PMID information were obtained from the GlyCosmos Portal (https://glycosmos.org; ref. 47) and GlyTouCan (https://glytoucan.org; ref. 21), using the data snapshot retrieved on 30 January 2026. Glycan-PMID associations were obtained from GlyGen (https://www.glygen.org; ref. 49) [GlyGen data release version: to be specified]. MeSH descriptor data were obtained from the 2026 release of Medical Subject Headings distributed by the U.S. National Library of Medicine (https://www.nlm.nih.gov/mesh). The reused experimental datasets analysed in this study are available from their original sources: the human macrophage glycomics data (ref. 33) [repository and accession, e.g.

PRIDE/MassIVE: PXD / MSV ], the human Alzheimer’s disease brain glycoproteomics data (ref. 34) [ProteomeXchange/PRIDE accession: PXD ], and the mouse brain glycoproteomics data (ref. 35) [ProteomeXchange/PRIDE accession: PXD ]. The GlycoMeSH-DB annotation database generated in this study, together with the trained GlycoMeSH-BERT model and the code snapshot, has been deposited in Zenodo under the DOI 10.5281/zenodo.20266489 and is also available through the GlycoMeSH-EA web tool. Source data for Figs. 2-5 are provided with this paper.

## Code availability

The source code used in this study is available at https://github.com/matsui-lab/GlycoMeSH-BERT_v1 under the CC-BY 4.0 license. A snapshot of the code, trained model, and GlycoMeSH-DB corresponding to the version used in this study has been archived in Zenodo with the DOI 10.5281/zenodo.20266489. The web-based enrichment analysis tool, GlycoMeSH-EA, is available at https://igcore.cloud/GlycoMeSH-EA/. A Docker image for local deployment of GlycoMeSH-EA is available from GitHub Container Registry at <u>ghcr.io/matsui-lab/glycomesh-ea:2026.1</u>. The GlycoMeSH-DB is available as downloadable files from the Zenodo archive and through GlycoMeSH-EA. GlycoMeSH-DB will be versioned by release year, for example GlycoMeSH-DB v2026.1, together with the corresponding MeSH release and GlyTouCan/GlyCosmos data snapshot. Database updates are planned annually, in accordance with the yearly MeSH term release schedule.

## Acknowledgements

This work was supported by the Human Glycome Atlas Project (HGA).

## Author contributions

A.K. conceived the study, curated the data, developed the methodology and software, performed the formal analyses, investigation, validation and visualization, and wrote the original draft. B.Z. and K.H. contributed to conceptualization, methodology, formal analysis, investigation and manuscript review. Y.M. acquired funding, administered the project, supervised the study and contributed to manuscript review and editing. All authors reviewed and approved the final manuscript.

## Ethics declarations

This study was based entirely on publicly available datasets and did not involve newly collected human or animal samples. Ethical approval, informed consent and animal experiment approval, where applicable, were obtained in the original studies.

## Supplementary information

