## Supplementary Information for "GlycoMeSH: linking glycan structures to biomedical context for systematic enrichment analysis"

### Supplementary Fig. 1: Literature-derived document-level MeSH contexts are informative but sparse across glycans.

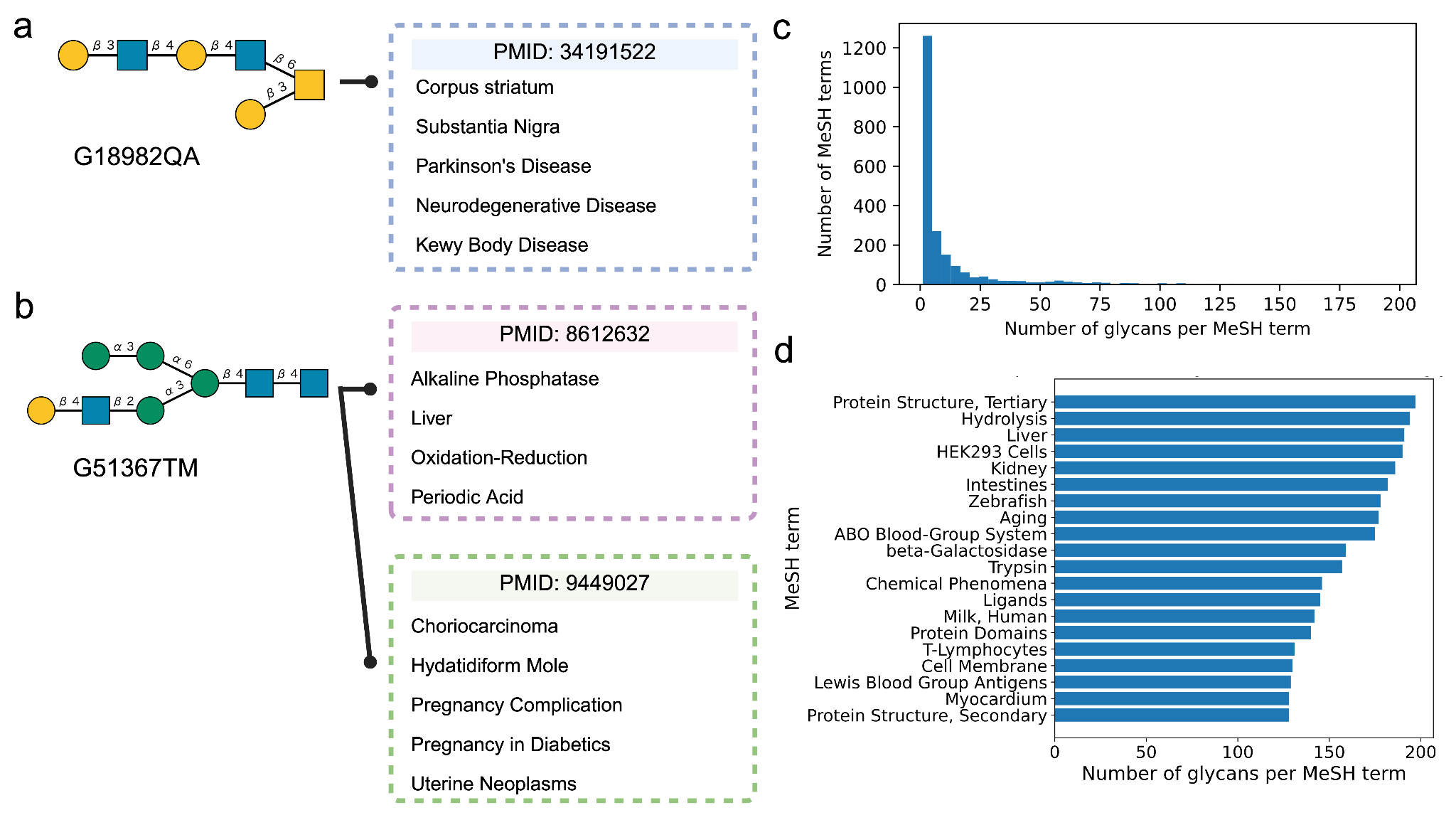

a,b, Examples of document-level MeSH contexts assigned to publications reporting glycans G18982QA (a) and G51367TM (b). The disease, anatomical, molecular and biological terms describe the source publications and do not necessarily denote direct glycan–function relations. c, Distribution of the number of glycans associated with each MeSH term in the curated literature-derived set. d, Top 20 MeSH terms by number of associated glycans.

### Supplementary Fig. 2: MeSH encoder selection and additional retrieval performance of GlycoMeSH-BERT

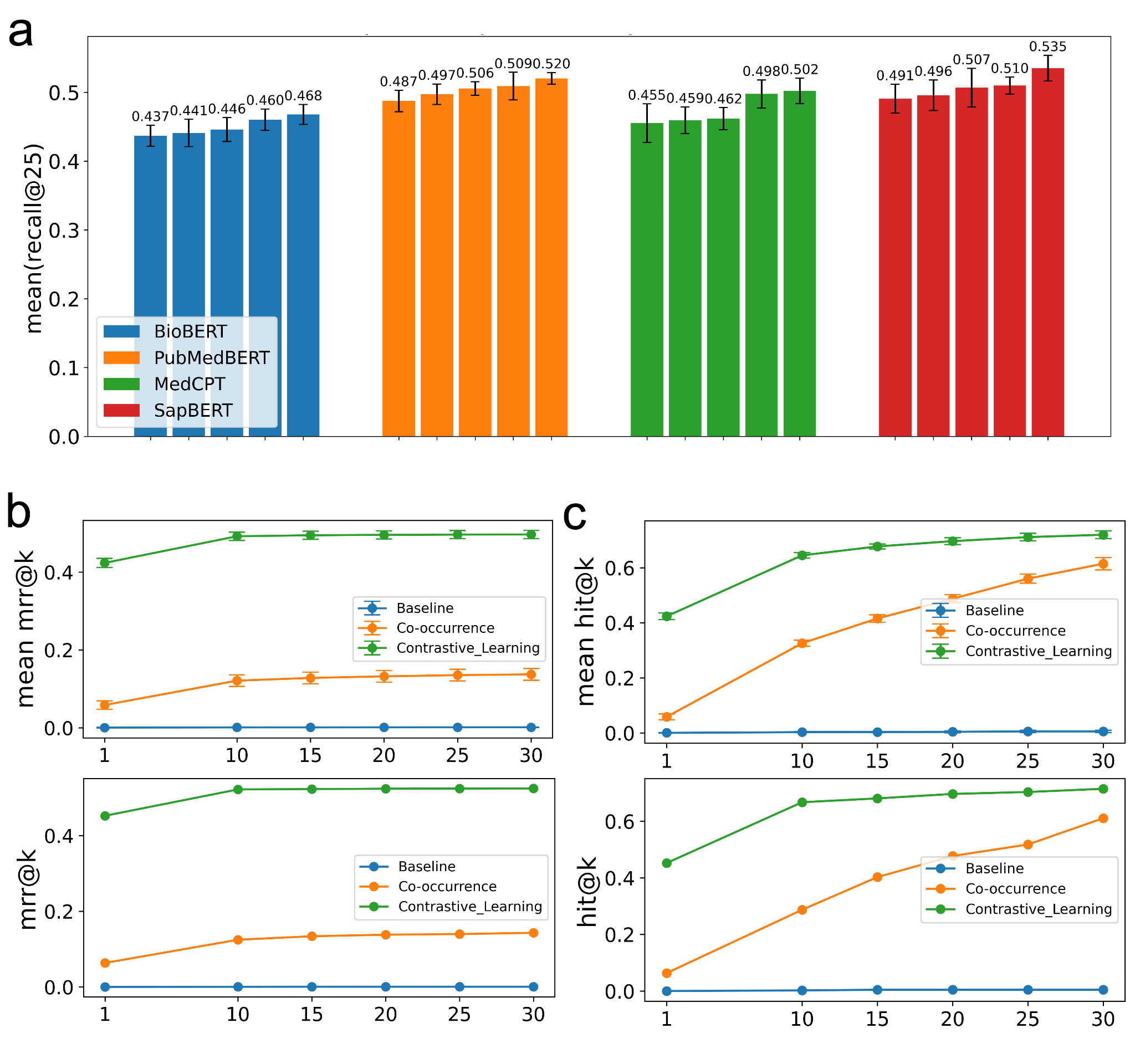

**a**, Performance of different biomedical language models evaluated by random hyperparameter search. The top five hyperparameter configurations for each model are shown, ranked by mean recall@25 across five-fold cross-validation; bars indicate mean ± s.d. **b**,**c**, Performance of GlycoMeSH-BERT evaluated using MRR@k (b) and hit@k (c). Top, cross-validation evaluation. Bottom, test dataset.

### Supplementary Fig. 3: Model performance stratified by similarity between test glycans and the training set

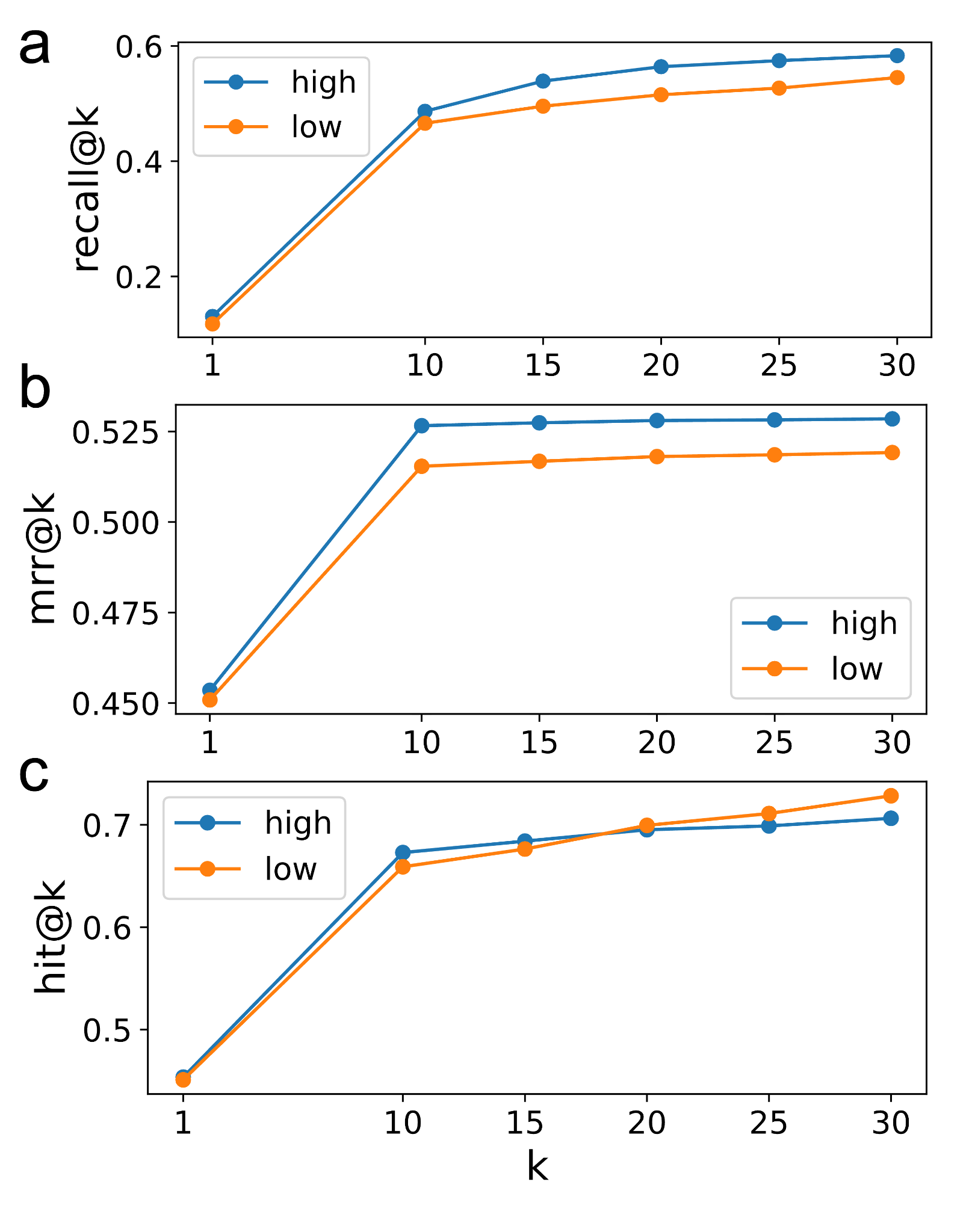

Test glycans were stratified based on their cosine similarity to glycans in the training set, and prediction performance was evaluated for each group. **a**, Recall@k. **b**, Mean reciprocal rank (MRR@k). **c**, Hit rate (hit@k).

### Supplementary Fig. 4: Comparison of prediction confidence and annotation coverage between recall@10 groups

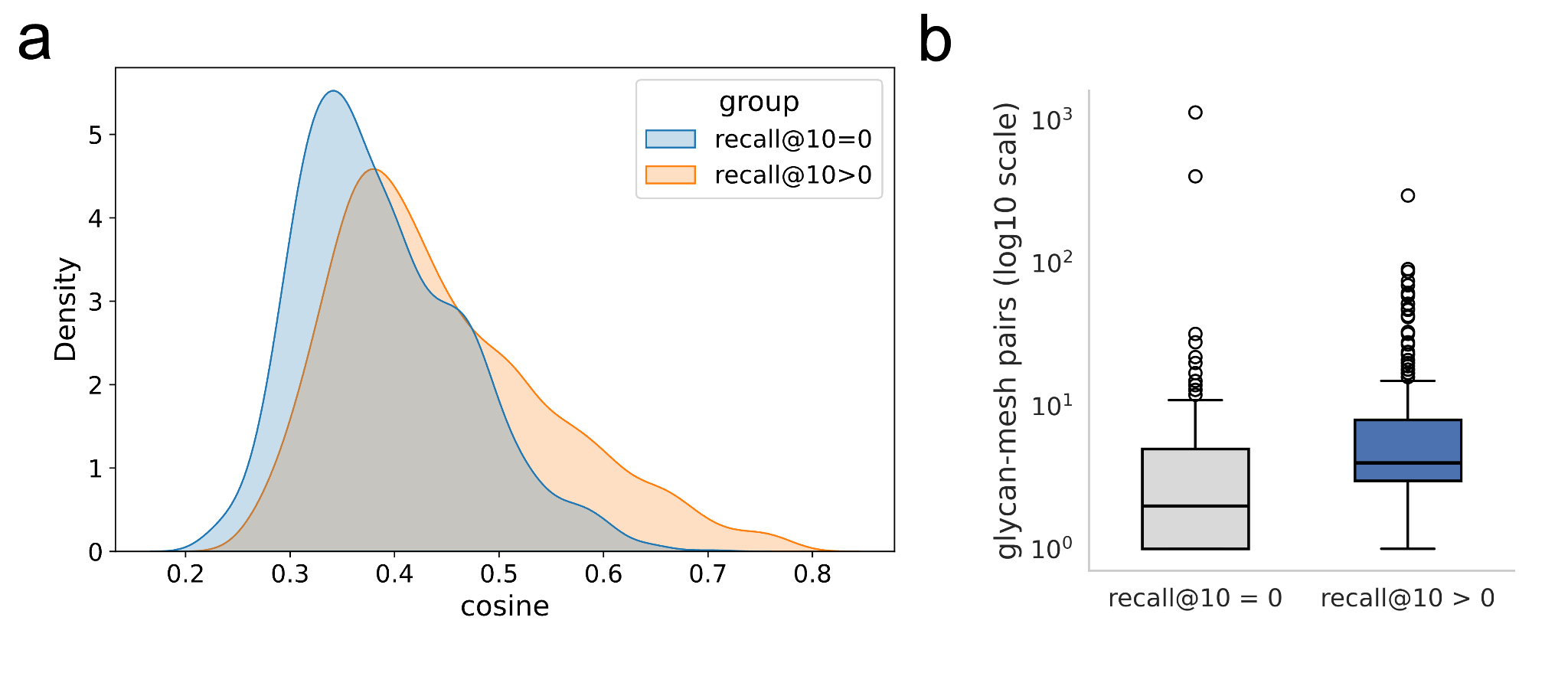

**a**, Distribution of cosine similarity scores for the top 10 predicted MeSH terms, comparing glycans with recall@10 = 0 and recall@10 > 0. **b**, Number of associated MeSH labels per glycan in each group. Statistical significance was assessed using the Mann–Whitney U test.

### Supplementary Fig. 5: Non-contrastive baseline architectures and annotation-space diversity.

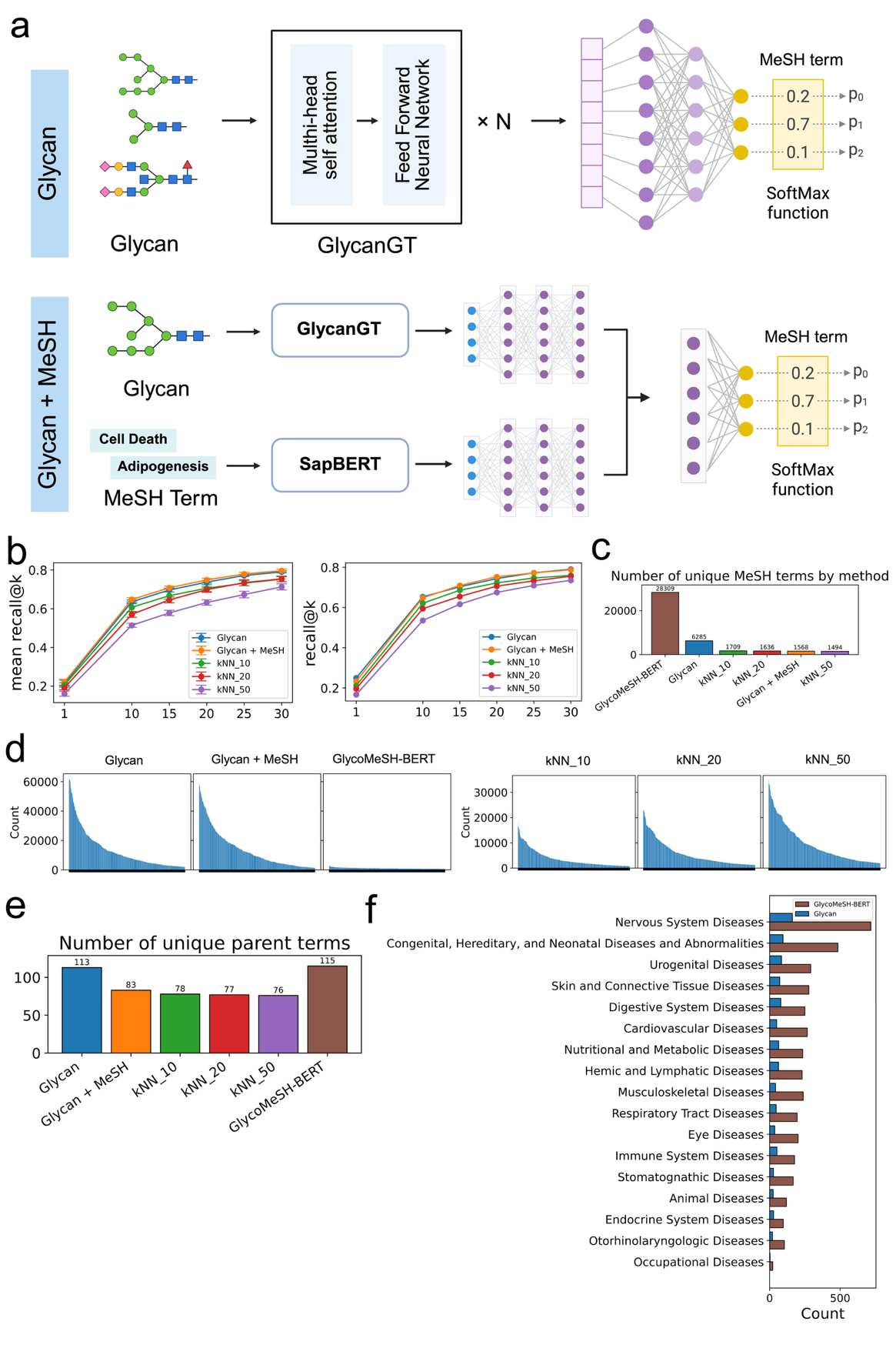

a, Model architectures of the MLP-based approaches. The glycan-only model takes GlycanGT-derived glycan embeddings as input, whereas the glycan+MeSH model incorporates both glycan and SapBERT-derived MeSH embeddings. b, Recall@k. Left, five-fold cross-validation mean ± s.d.; right, performance on the held-out glycan test set. c, Number of unique MeSH terms among the aggregated top 100 predictions. d, Frequency distribution of MeSH terms in those predictions. e, Number of represented MeSH parent terms. f, Term-level diversity within Disease-related parent categories for GlycoMeSH-BERT and the glycan-only MLP.

### Supplementary Fig. 6: Threshold sensitivity analysis for GlycoMeSH-DB construction

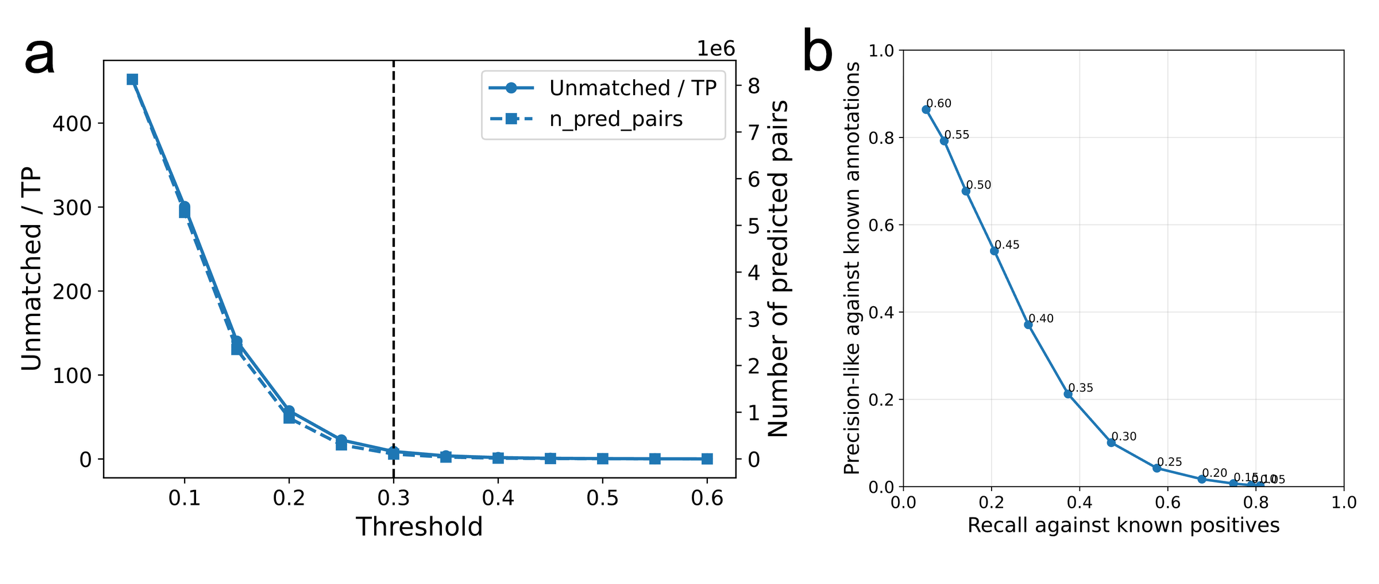

a, Trade-off across cosine-similarity thresholds between the number of predicted glycan–MeSH associations and the ratio of predictions absent from the current literature-derived set to matched literature-derived pairs (Unmatched/TP). b, Retention-versus-expansion summary across thresholds. Recall-like values denote retention of current literature-derived pairs. Unmatched predictions denote absence from the incomplete curation, not validated false positives; the plotted precision-like quantity is therefore an operational threshold summary rather than a precision estimate or validation of individual associations.

### Supplementary Fig. 7: Illustrative application of GlycoMeSH-EA to glycoproteomics data from an Alzheimer’s disease mouse model.

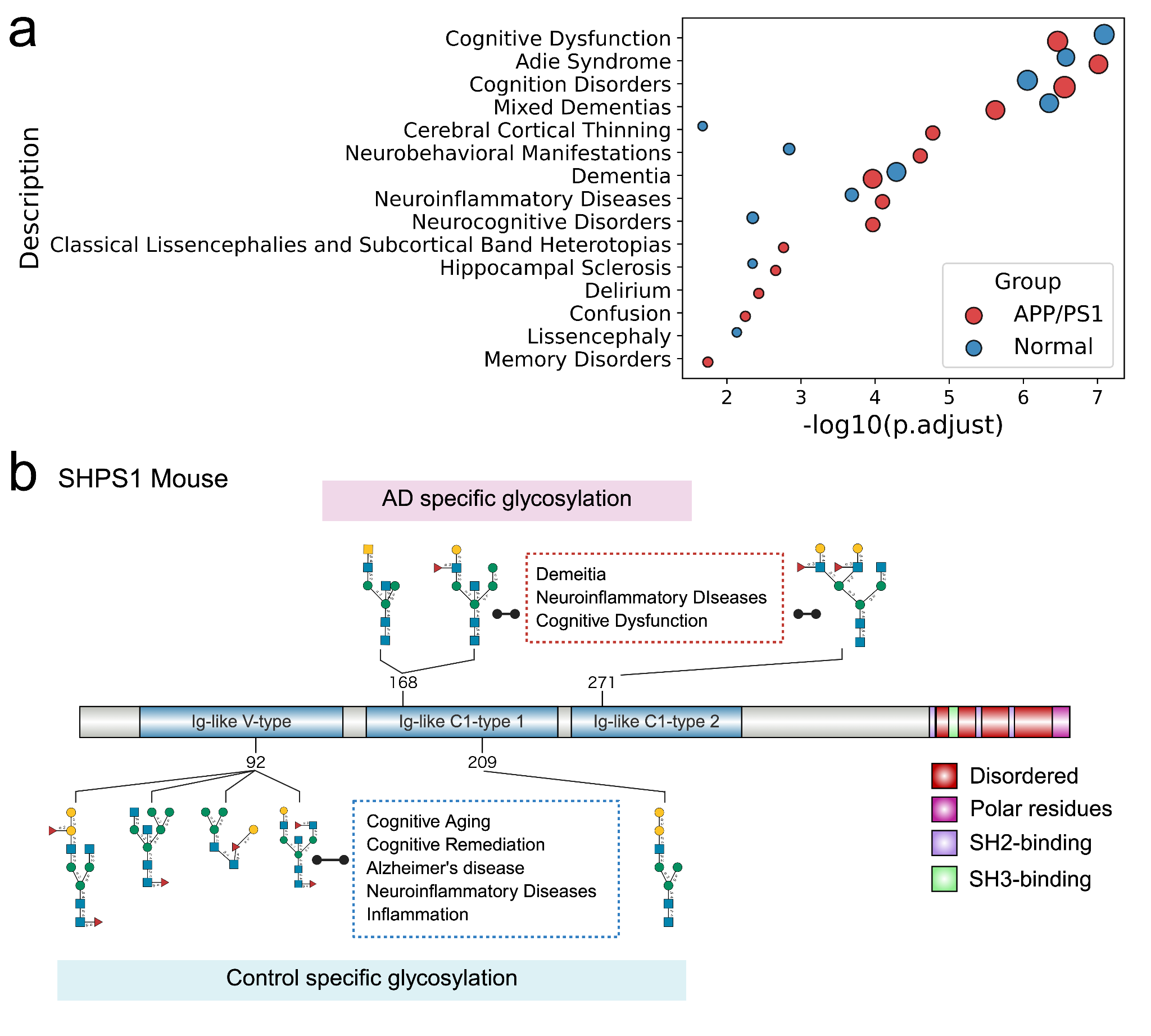

a, GlycoMeSH-EA results for glycans detected in APP/PS1 and control mouse samples, shown for the pre-selected display categories. ORA used the hypergeometric test (enricher, clusterProfiler), all glycans in the analysis as background and Benjamini–Hochberg correction across the full MeSH vocabulary. The nervous-system categories were selected in advance for display only, and unrestricted results are provided in Supplementary Table 18; the panel illustrates method use and is not an independent validation of disease specificity. b, Descriptive SHPS1 domain-level glycosylation patterns. SHPS1 was selected post hoc as a hypothesis-generating example, and the condition-specific glycans shown were not statistically tested for domain-localization specificity.
